# Psilocybin biphasically modulates cortical and behavioral activity in mice

**DOI:** 10.1101/2024.01.18.576229

**Authors:** Adam T. Brockett, Nikolas A. Francis

## Abstract

Psilocybin is a serotonergic psychedelic believed to have therapeutic potential for neuropsychiatric conditions. Despite well-documented prevalence of perceptual alterations, hallucinations, and synesthesia associated with psychedelic experiences, little is known about how psilocybin affects sensory cortex or alters the activity of neurons in awake animals. To investigate, we conducted 2-photon imaging experiments in auditory cortex of awake mice and video analysis of mouse behavior, both at baseline and during psilocybin treatment. We found biphasic effects of psilocybin on behavioral and cortical activity. A 2 mg/kg dose of psilocybin initially increased behavioral activity and neural responses to sound. 30 minutes post-dose, mice became behaviorally hypoactive and cortical responses to sound decreased, while neural response variance and noise correlations increased. In contrast, neuronal selectivity for auditory stimuli remained stable during psilocybin treatment. Our results suggest that psilocybin modulates the role of intrinsic versus stimulus-driven activity in sensory cortex, while preserving fundamental sensory processing.

**Graphical Abstract.**
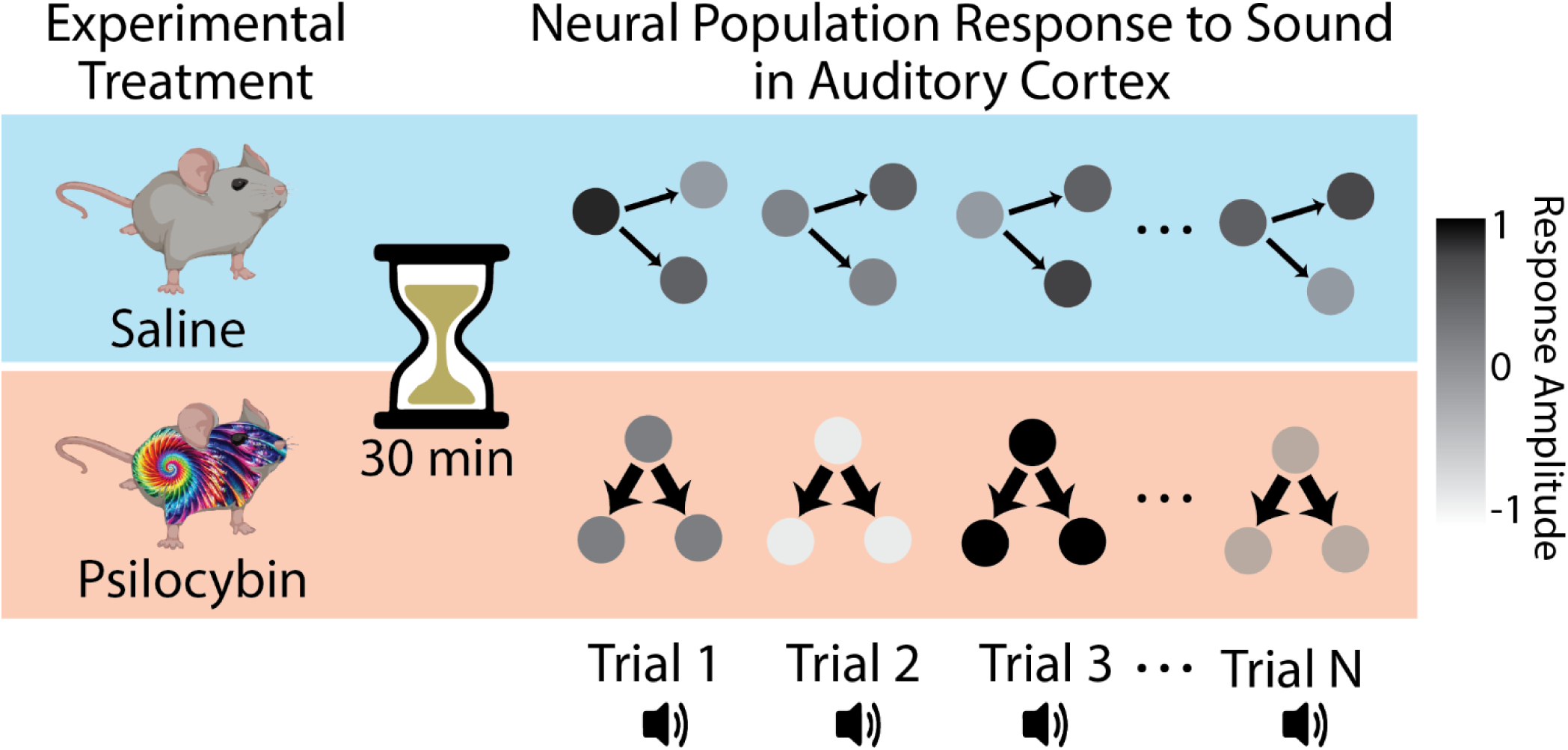
Summary of psilocybin’s effect on auditory cortical responses to sound in mice. 30 minutes after injecting the inert vehicle saline, typical auditory responses occur within a relatively narrow range of possible amplitudes, i.e., each neuron’s response variance is weakly correlated with a neighboring neuron’s response variance. In contrast, 30 minutes after injecting 2 mg/kg of psilocybin, population response variance becomes more correlated between individual neurons, and the range of response amplitudes increases. These findings suggest that psilocybin modulates the role of intrinsic versus stimulus-driven neural activity in sensory perception, which may serve as a basis for auditory hallucination at the level of neuronal micro-circuits.

## INTRODUCTION

Psilocybin is a psychoactive prodrug long known to induce atypical changes in conscious experience including altered perception, cognition, and mood. The therapeutic potential of psilocybin, and other serotonergic psychedelics, has been studied off and on for well over fifty years^1–4^. Recently, clinical interest in psilocybin has increased following several small clinical trials showing that 1-3 treatments with psilocybin can produce lasting reductions in the severity of depressive symptoms^5–10^. Despite these promising results, few studies have investigated the cellular and physiological mechanisms by which psilocybin produces these profound changes in brain state.

Clarifying how psilocybin affects cortical neurophysiology is essential for psilocybin’s continued emergence as a safe and viable treatment option. A recent study in mice found that psilocybin increased neuronal spiking in anterior cingulate cortex (ACC), while modulating synchrony in local field potentials^11^. Functional magnetic resonance imaging (fMRI) studies in humans and rats have shown that psilocybin alters brain-wide activity and synchronization^10,12,13^, and that altered perceptions associated with psilocybin can be suppressed via blockage of the 5-HT2a receptor^14,15^. However, it remains unclear how psilocybin’s effects on cortical activity produce the perceptual alterations, hallucinations, and synesthesia that may occur during psychedelic experiences^14,16^.

Here, we used 2-photon (2P) Ca^2+^ imaging^17^ of neural activity in awake mice to study how psilocybin affects sensory processing in primary auditory cortex (A1) layer 2/3 (L2/3). We also used video analysis of free-roaming mouse behavior to track head-twitch responses (HTRs) and overall movement around a cage. We show that psilocybin biphasically modulates both cortical and behavioral states, while preserving neural selectivity for auditory stimuli. Importantly, our results suggest that increased auditory response variance and functional connectivity in A1 L2/3 during psilocybin treatment may reflect perceptual alterations or pseudo-hallucinations.

## RESULTS

### Psilocybin has a biphasic effect on auditory cortex

Since auditory cortex is critical for auditory perception and is thought to play a role in auditory hallucination^18^, we sought to clarify the effects of psilocybin on the auditory responsiveness of individual neurons in A1 L2/3 using 2P imaging (*see* STAR Methods). Importantly, all experiments were done on awake mice.

Figure 1a illustrates the timeline of our imaging experiments. For each mouse, we began with a pre-injection imaging session (Pre). We then waited an hour before an intraperitoneal (IP) injection of either a 2 mg/kg dose of psilocybin (N=6 mice, N=8 experiments, dissolved in 0.1 ml saline) or the 0.1 ml saline vehicle alone (N=3 mice, N=6 experiments). The 2 mg/kg dose was chosen to approximate doses used in human clinical trials^5–10^. Psilocybin was provided by the NIDA Drug Supply Program.

**Figure 1.**
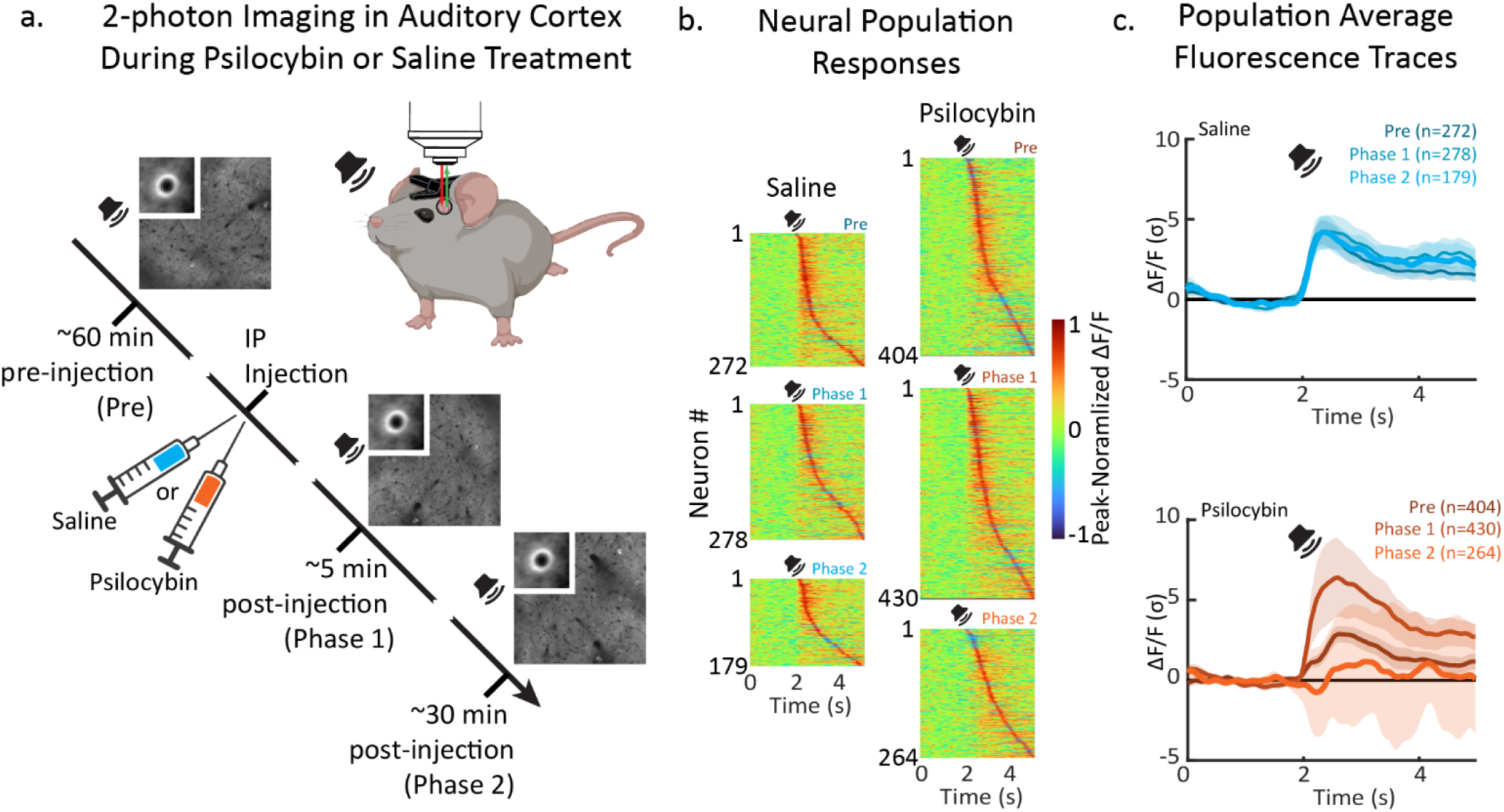
Psilocybin biphasically modulates neural responses to sound in primary auditory cortex (A1) layer 2/3 (L2/3) of awake mice. **a. Experimental paradigm.** Mice were imaged pre-injection (Pre), then given an intraperitoneal (IP) injection with either 2 mg/kg psilocybin or 0.1 ml saline, then two post-injection imaging sessions occurred (Phase 1 and Phase 2, respectively). Color-coding here for psilocybin (orange) vs saline (cyan) is used for all remaining panels. Randomized pure-tones were presented during imaging. **b. Imaged Neural Population.** The auditory responses of the imaged population from each experimental phase and condition are shown as heatmaps. Hot and cool colors show peak-normalized fluorescence (ΔF/F) for each imaged neuron. **c. Population-averaged auditory responses in A1 L2/3.** The darkest, lighter, and lightest colored lines show responses during Pre, Phase 1 and Phase 2 sessions, respectively. Shading shows 2 standard errors of the mean (SEM).

The injected mouse was placed back under the microscope for post-injection imaging (Phase 1), which typically began within 5 minutes after injection. With the injected mouse still under the microscope, we then performed a final imaging session (Phase 2), which typically began within 30 minutes after injection. Each imaging session occurred in a different 2P field of view and lasted approximately 20 minutes while we presented 70 dB SPL, 0.5 s pure-tones at 10 frequencies (2-45 kHz) from a free-field speaker. Each frequency was repeated 20 times in randomized order. Stimuli were presented at randomized inter-trial intervals of 6, 7, or 8 s.

Figure 1b shows heatmaps of the populations of auditory-responsive neurons that were recorded for each experimental condition. Each row of each heatmap shows an individual neuron’s average activity during a trial of the experiment. Most neurons began each trial in a quiescent state for 2 seconds, followed by a sudden rise in activity after stimulus presentation. As expected, auditory responses were similar across Pre, Phase 1, and Phase 2 sessions for mice injected with saline (figure 1c, top panel; figure 2a). In contrast, psilocybin induced a biphasic modulation of neural responses to sound (figure 1c, bottom panel; figure 2a,b). Immediately after injection during psilocybin Phase 1, neurons in A1 L2/3 became hyper-responsive to sound, relative to the psilocybin Pre session (figure 2a; p=0.006). In contrast, 30 minutes post-injection during psilocybin Phase 2, the average neural response amplitude decreased relative to psilocybin Phase 1 (figure 2a; p=0.015).

**Figure 2.**
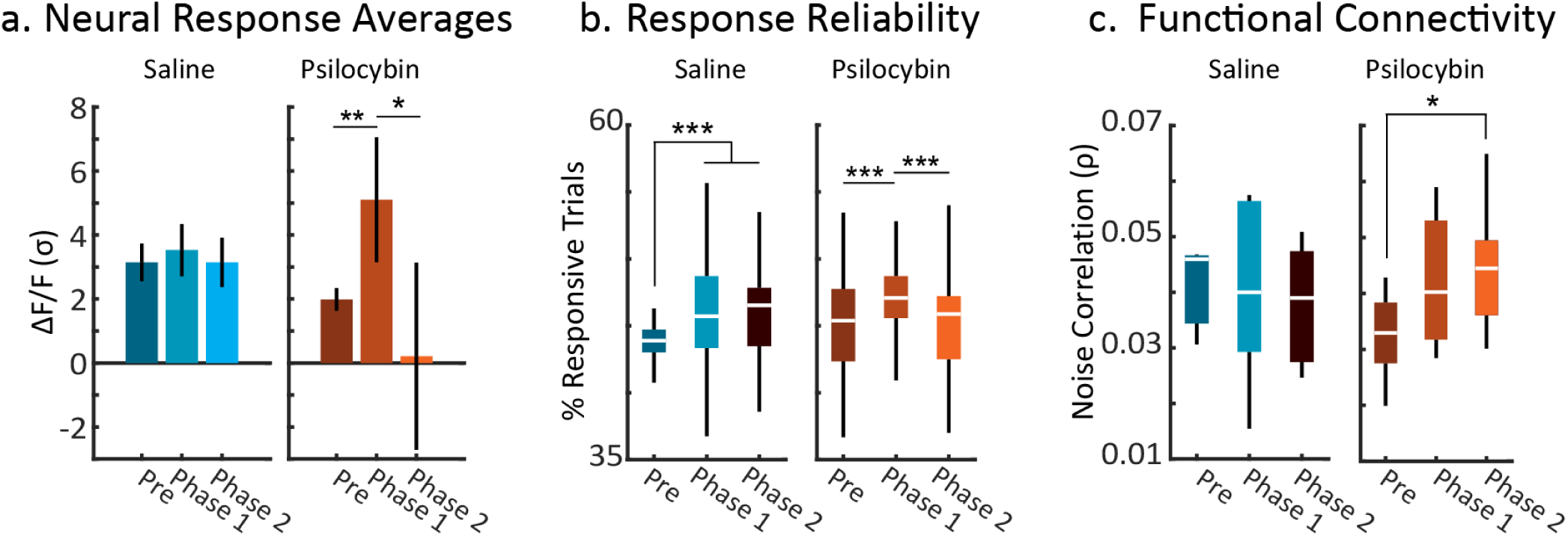
Psilocybin modulates neuronal response variability, reliability, and functional connectivity in A1 L2/3. **a. Time-averaged neural responses.** Activity during the 1 s following stimulus presentation was averaged for each neuron. The mean population response for each condition is shown with 2 SEMs. The stars indicate significant differences in the Psilocybin condition (Pre vs Phase 1: p=0.006; Phase 2 vs Phase 1: p=0.015). **b. Response Reliability.** Neuronal response reliability was quantified as the percent of pure-tone responsive trials for each condition. Stars indicate significant differences (Saline: Pre vs Phase 1 and Phase 2, p<0.001; Psilocybin: Pre vs Phase 1, p<0.001; Phase 1 vs Phase 2, p<0.001). **c. Functional connectivity.** Noise correlations were computed for pairs of neurons in each 2-photon field of view. Each panel in figures 2b and 2c show a box plot for the data in each condition. The boxes show interquartile ranges, white lines show median values, and whiskers show the range of the distribution. The star in figure 2c indicates a significant Pre vs Phase 2 difference in noise correlations (p=0.045).

To determine if psilocybin changes how often neurons in auditory cortex respond to sound, we measured neural response reliability (figure 2b). For a given neuron, response reliability is the percentage of trials in which the activity immediately following an auditory stimulus exceeds the activity during the 2 s of silence preceding the stimulus. Thus, a 50% response reliability would describe a neuron that had an auditory response for 10/20 trials. We found that response reliability tended to increase during saline treatment (Pre vs Phase 1: p<0.001; Pre vs Phase 2: p<0.001), but was similar for Phases 1 and 2 (p>0.05). It may be that stress related to IP injection *per se*, or simply repeating the stimulus set many times across phases, enhances response reliability. During psilocybin treatment, response reliability increased during Phase 1 (Pre vs Phase 1: p<0.001), similar to saline treatment. However, unlike saline, response reliability then decreased back to baseline levels during psilocybin Phase 2 (figure 2b; Pre vs Phase 2: p>0.05; Phase 1 vs Phase 2: p<0.001). In other words, auditory responses became slightly less reliable during psilocybin Phase 2. Thus, psilocybin has a biphasic effect on both the likelihood that a neuron will respond to sound and the size of the neuron’s response to sound.

### Psilocybin increases neuronal response variability and functional connectivity in auditory cortex

It is important to note that during psilocybin Phase 2, while the average neural response amplitude decreased, neural response *variance increased* (see shading in figure 1c, bottom panel; see error bars in the right-most bar graph in figure 2a). This means that the amplitude of responses to sound spanned a wider range during psilocybin treatment. To investigate whether the increased response variance was patterned, we computed trial-to-trial correlations of response variance between pairs of neurons^19–21^, i.e., ‘noise correlations’. An increase in noise correlation indicates an increase in functional connectivity between pairs of neurons^19–21^.

We measured noise correlations during Pre, Phase 1, and Phase 2 time periods for both saline and psilocybin treatments (figure 2c). We found that noise correlations tended to remain constant during Pre, Phase 1, and Phase 2 of saline treatments (left panel, figure 2c; p>0.05). In contrast, noise correlations significantly increased during psilocybin Phase 2, compared to Pre (right panel, figure 2c; p=0.043) (figure 2c). This increase in noise correlation, i.e., the increase in correlated response variance between pairs of neurons, indicates that psilocybin increased functional connectivity in A1 during the same period when mice transitioned into a more hypoactive behavioral state. Importantly, noise correlations are unbiased by stimulus-driven neural responses. Thus, our results suggest that psilocybin enhances the role of intrinsic brain influences on sensory processing in A1 L2/3.

### Psilocybin does not change frequency tuning in auditory cortex

So far, we have shown that psilocybin biphasically alters neuronal responses to sound in A1 L2/3. However, some aspects of auditory responsiveness in A1, such as pure-tone frequency selectivity, i.e., ‘frequency tuning’, are inherited from a frequency-specific labelled-line that originates in the cochlea, rises through the brainstem, and reaches the cortex via the medial geniculate body of the thalamus. Thus, neuronal frequency tuning curves (FTCs) might be expected to withstand the response amplitude changes induced by psilocybin. To examine this, we measured the FTC of each imaged neuron, and then grouped neurons by ‘Best Frequency’ (BF), i.e., the frequency corresponding to the FTC peak. We then peak-normalized each neuron’s FTC and averaged the FTCs for each BF. This produced 10 FTCs per experimental condition (figure 3a). We found that FTCs were well-tuned, i.e., selective for the BF, and were similarly shaped across all experimental conditions. We quantified changes in FTC shape by measuring the size of off-BF responses relative to BF (figure 3b). We found that the off-BF response magnitudes remained stable throughout experiments. Thus, psilocybin did not change frequency tuning in A1 L2/3.

**Figure 3.**
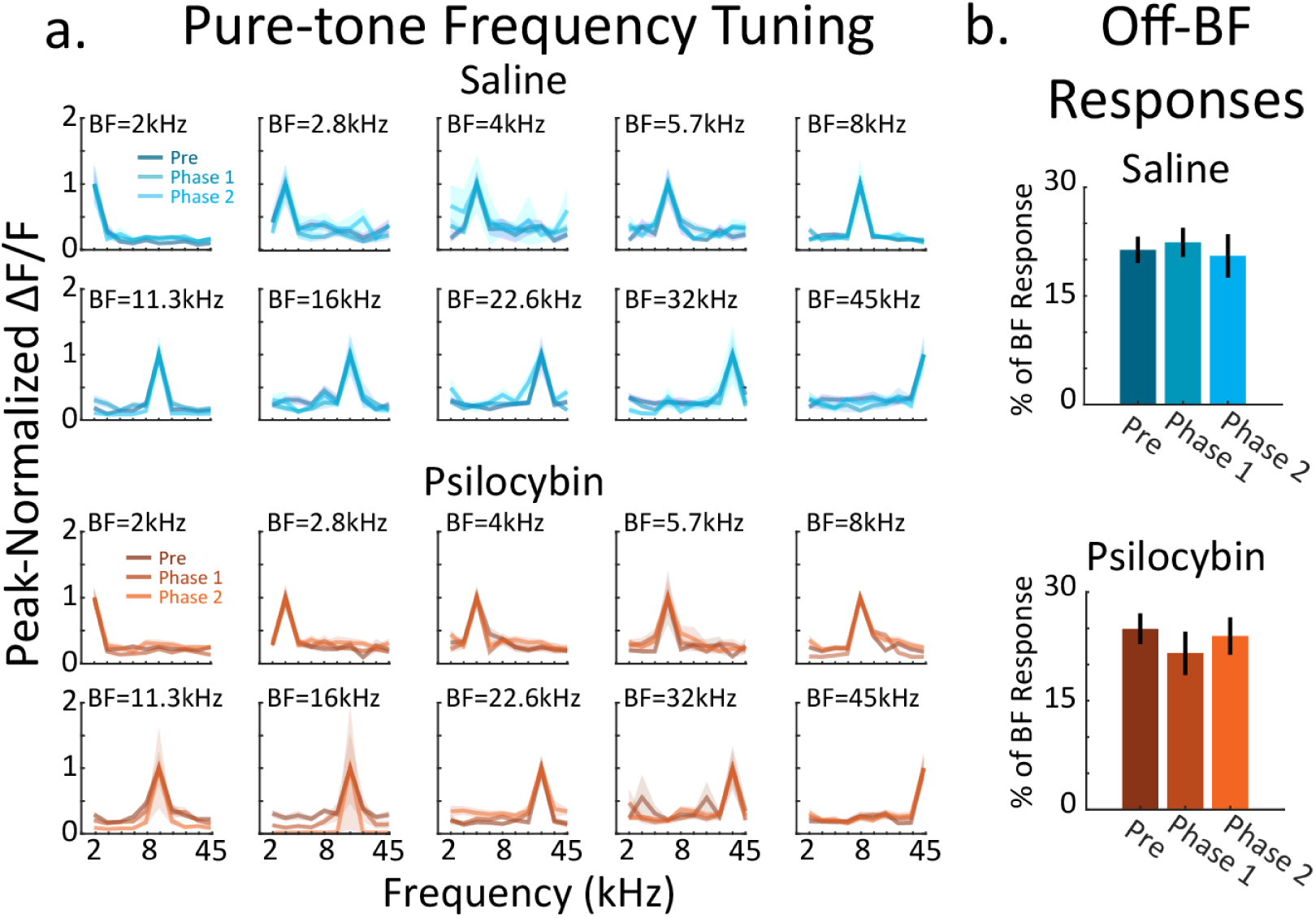
Psilocybin does not change frequency tuning in A1 L2/3. **a.** Frequency tuning curves (FTCs) were computed for each imaged neuron by finding the average response to each of the 10 pure-tone frequencies (2-45 kHz) presented at 70 dB SPL. The best frequency (BF), i.e., the frequency with the biggest response, was found for each neuron, and then FTCs from neurons with the same BF were averaged. The FTCs from each condition are shown in panel a. Each panel contains three overlapping FTCs. Shading shows 1 SEM. **b. Average neural responses to off-BF frequencies.** No change in FTC selectivity was found across conditions. FTC selectivity was quantified as the average response to pure-tones at frequencies other than the BF of a given neuron. Responses are reported as the percent of the response at BF.

### Psilocybin has a biphasic effect on mouse behavior

Having established a biphasic time-course for psilocybin’s effect on auditory cortex, we hypothesized that a similar biphasic modulation of behavior may occur. To quantify psilocybin’s effects on the behavior of mice, we built a custom arena for video tracking of mouse movement (*see* STAR Methods). The arena consisted of a large black acrylic box that contained a smaller clear acrylic box in the center of the arena. Videos were taken at 150 frames per second (fps) and analyzed at 30 fps (*see* supplementary videos S1 and S2). Experiments began with habituation, by placing the mouse in the small clear box for an hour, before administering an IP injection of either psilocybin (2 mg/kg in 0.1 ml saline) (n=5 mice) or 0.1 ml saline vehicle (n=5 mice) (figure 4a). Immediately after injection, the mouse was placed back in the clear box, covered with a clear lid, and then videoed for an additional 40 minutes (figure 4b).

**Figure 4.**
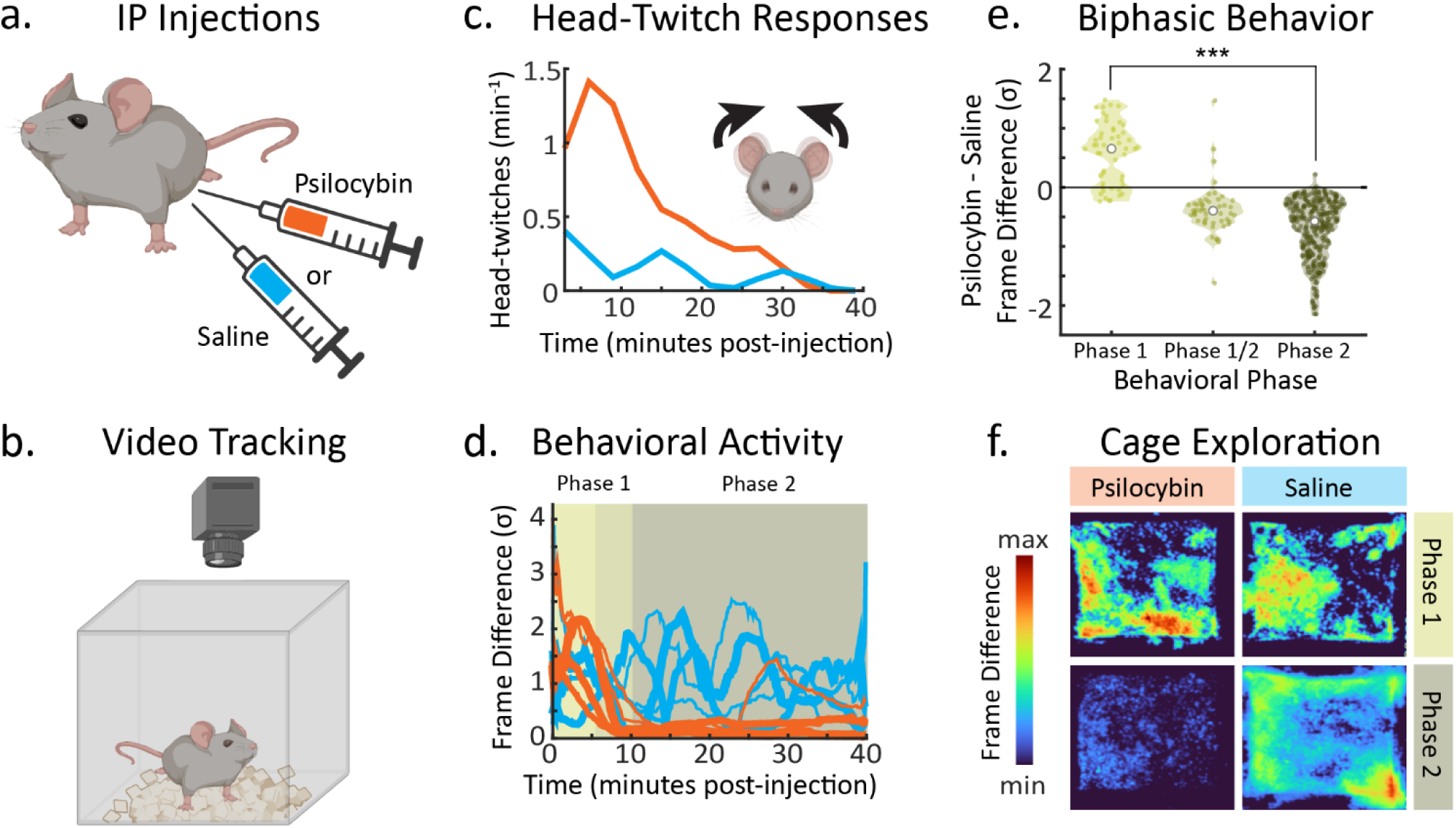
Psilocybin increases head-twitch responses and biphasically modulates overall mouse movement. **a. Experimental preparation.** Mice were given an IP injection of either saline (cyan) or 2 mg/kg psilocybin (orange). The color-coding used here for experiments with psilocybin vs saline is also used in panels c, d, and f. **b.** Injected mice were placed inside a video recording cage and their behavior was recorded for 40 minutes at 150 frames per second (fps). **c. Time-course of head-twitch responses.** Head-twitches were manually scored in 2 mice (n=1 psilocybin; n=1 saline). **d. Time-course of overall mouse movement.** The magnitude of mouse movement was quantified on a frame-by-frame basis and plotted for each treatment (psilocybin: n=5; saline: n=5). **e. Biphasic behavioral effects of psilocybin.** Frame-wise differences from panel d are shown as violin plots for phase 1 and 2, color coded as in panel d. The white dots show median values for each distribution. The stars show that the means of Phase 1a and 2 were significantly different (p<0.001)**. f. Cage exploration.** The magnitude of mouse movement across space in the cage was averaged across frames for each mouse. Examples for psilocybin (left) and saline (right) injected mice are shown here. Movement was averaged in a time-window corresponding to psilocybin effect phase 1 (0-10 minutes post-injection) and phase 2 (10-40 minutes post-injection). Hot and cool colors indicate regions with a high and low magnitude movement, respectively.

Previous research on the behavioral effects of psilocybin measured HTRs as a dose-dependent indicator of psychedelic drug effects^22,23^. Here, we successfully replicated previous findings of the HTR time-course by manually labeling video for each occurrence of a HTR, after both psilocybin (n=1 mouse) and saline injections (n=1 mouse; figure 4c, supplemental videos S1 and S2). Consistent with previous work^22^, we found that the rate of head-twitches was greater for psilocybin than saline, increased quickly after psilocybin injection, and then declined within 10 minutes.

During behavioral labelling, we noticed that once HTRs declined, mice injected with psilocybin became very still within the clear box. Thus, we decided to compare overall behavioral movement across each video for mice injected with psilocybin versus saline (figure 4d, supplemental video S2; *see* STAR Methods for details). Figure 4d shows the movement traces for both saline and psilocybin injected mice. The data show clear time-dependent differences between the movements of psilocybin versus saline mice, which we tested for significance as a function of time using bootstrap t-tests in a 5-minute sliding window (50% overlap).

We identified a biphasic behavioral effect of psilocybin that mirrored its effect on cortical responses to sound, with no significant difference compared to saline only in the 7.5-10 minute post-injection range. Figure 4e shows the mouse movement distributions for each phase of behavioral changes. In Phase 1 (0-10 minutes), the mice injected with psilocybin became hyperactive, relative to mice injected with saline. This period contains the rise in HTRs (figure 4c). In Phase 2 (10-40 minutes post-injection), psilocybin mice became hypoactive relative to controls. We separately analyzed differences in the 7.5-10 minutes transition period between phases 1 and 2 and found that while the median effect resembled phase 2, the variance of differences spanned the range of both phase 2 and early phase 1. Thus, 10 minutes marks an approximate time-point in which the effects of psilocybin on behavior switch from more stimulant to more sedative in nature.

To gain further insight into the effects of psilocybin on mouse behavior, we also quantified cage exploration using frame differences as described above, while preserving spatial information. Figure 4f shows spatial movement for psilocybin versus controls in behavioral phases 1 and 2. The results show a similar pattern as described above, where saline mice continued to explore the cage throughout the experiment, whereas psilocybin mice ceased exploration 10 minutes post-injection. Together, the time-courses of HTRs, the magnitude of behavioral movements, and the scope of cage exploration describe a similar biphasic time-course of behavior following psilocybin treatment: a transient period of hyperactivity followed by the sudden transition into a longer period of hypoactivity. This time-course mirrors our findings on the effects of psilocybin in auditory cortex.

## DISCUSSION

Our results show how individual neurons in sensory cortex of awake mice are biphasically modulated by psilocybin, the prodrug found in mushrooms of the genus *Psilocybe,* which have been ingested by humans for thousands of years and show promising therapeutic potential for neuropsychiatric conditions. However, our understanding of the cellular effects of psilocybin in the brain, and serotonergic psychedelics more generally, is limited. Here, we show that shortly after treating mice with a 2 mg/kg dose of psilocybin, there is an increase in locomotion and neural responsiveness to sound. However, 30-minutes later this initial increase in behavioral and neural responsiveness sharply diminishes. Both behavioral and neural measures suggest that mice become less sensitive to external sensory cues, shown here as diminished HTRs, decreased locomotor exploration, and a suppression of average neural responsiveness to sound. Interestingly, while the average neural response to sound decreased, noise correlations increased, suggesting that local functional connectivity in A1 L2/3 increased during the behaviorally quiescent psychedelic effects.

Few studies have examined the response properties of individual neurons within the context of psilocybin treatment. Recently, it was shown that a 2 mg/kg dose of psilocybin in mice increased spiking and decreased the power of low frequency oscillations in the ACC, a frontal brain area important for cognitive control^11^. The decreased power was interpreted as a desynchronization of local neural activity. These findings are partially corroborated by a combined fMRI-immunofluorescence study in rats, which reported that a 2 mg/kg dose of psilocybin increased bold oxygen level dependent responding and functional connectivity in numerous frontal, parietal, and temporal lobe areas, as well as the striatum^13^. While neither study probed A1 directly, nor examined sensory processing in general, the reported psilocybin-induced modulation of local cortical synchrony mimics the changes we find here in A1.

Our examination of the response properties of neurons in A1 revealed a clear biphasic change in activity that preceded the increase in functional connectivity. This switch from a stimulus-driven state to a more intrinsically-driven state may reflect a change in the relative levels of top-down vs bottom-up control of A1. Recent work has shown that the orbitofrontal cortex (OFC) and ACC modulate auditory processing in A1^24–27^. Thus, our observed changes in A1 might arise due to concurrent effects of psilocybin on activity and synchrony in frontal regions like OFC and ACC and may contribute to auditory alteration or pseudo-hallucination. It is important to note that the fundamental auditory property of frequency tuning remained stable despite the other dynamic effects of psilocybin, which speaks to the limits in which hearing can be altered during a psychedelic experience. Future work should investigate differential influences of bottom-up vs top-down control of sensory cortex during serotonergic psychedelic treatment.

In summary, our findings speak to the role of psilocybin in modulating the neural basis of sensory perception. While psilocybin shows great promise as a therapeutic adjuvant, it remains unclear how psilocybin produces its therapeutic effects. Our findings outline how psilocybin affects the activity of individual neurons in an awake mammal, paving the way for *in vivo* investigation of psychedelic effects on cortical micro-circuits.

## Acknowledgements

This research was funded by the University of Maryland (UMD) Individual Grand Challenges Award (GC30) awarded to ATB, and a UMD Brain and Behavior Institute seed grant awarded to NAF. The authors thank Franshesca Orellana Castellanos, Jonathan Dinh, Gabrielle Stephens, and Sofia Leusch for labeling behavioral videos. The authors would like to thank the NIH NIDA Drug Supply Program for providing the psilocybin used in these experiments.

## Author Contributions

ATB and NAF designed the experiments. NAF designed experimental software and hardware, performed experiments, and analyzed the data. ATB and NAF wrote the manuscript.

## Declaration of Interests

The authors declare no competing interests.

## STAR METHODS

### RESOURCE AVAILABILITY

#### Lead contact

Further information and requests for resources should be directed to and will be fulfilled by the lead contact, Dr. Nikolas Francis

#### Material Availability

This study did not generate new unique reagents.

#### Data and code availability

Imaging data have been deposited in the Digital Repository at the University of Maryland and are publicly available as of the date of publication. DOIs are listed in the key resources table.

### EXPERIMENTAL MODEL AND SUBJECT DETAILS

All procedures were approved by the University of Maryland Institutional Animal Care and Use Committee. Psilocybin was procured from the National institute on Drug Abuse drug supply program. We used N=19 mice, which were F1 offspring of CBA/CaJ mice (The Jackson Laboratory; stock #000654) crossed with transgenic C57BL/6J-Tg(Thy1GCaMP6s)GP4.3Dkim/J mice^17,28,29^ (The Jackson Laboratory; stock #024275), 1.5-7 months old. We used the F1 generation of the crossed mice because they have healthy hearing at least 1 year into adulthood^28^. Mice were housed under a reversed 12 h-light/12 h-dark light cycle.

### METHOD DETAILS

#### Video analysis of mouse behavior

We studied mouse behavior using a custom-built video tracking arena. The arena consisted of a large black acrylic box with an open top, that contained a smaller clear acrylic box in the center of the arena. White LED arrays were attached to inside walls of the black box to illuminate the small clear box. Bedding was placed inside the clear box, and a pco.Panda 4.2 CMOS camera was mounted above, with only the extent of clear box kept in-frame. Videos were taken at 150 frames per second (fps) and analyzed at 30 fps (*see* supplementary videos S1 and S2). We quantified mouse movement in each video, V, by first creating a new video, V’, from sequential frame differences in V, i.e., V’(f) = V(f+1)-V(f), where f indexes each frame. We then created a 1-dimensional frame difference trace, D, by summing over all pixels for each frame in V’. We extracted the envelope of D, using its low-passed Hilbert transform magnitude, to produce the mouse’s movement trace, M, spanning each 40-minute video.

Behavioral experiments began with habituation, by placing the mouse in the small clear box for an hour, before being administered an intraperitoneal (IP) injection of either psilocybin (2 mg/kg in 0.1 ml saline) (n=5 mice) or a 0.1 ml saline vehicle (n=5 mice) (figure 4a). Immediately after injection, the mouse was placed back in the clear box, covered with a clear lid, and then videoed for an additional 40 minutes (figure 4b). The 2 mg/kg dose was chosen to approximate the dose use in human clinical trials^5–10^.

#### Stimulus design and presentation

During imaging experiments, we presented awake mice with 70 dB SPL pure-tones from an ES1 free-field speaker (Tucker-Davis Technologies) connected to an ED1 amplifier (Tucker-Davis Technologies). The speaker was calibrated to give a flat frequency response in the 1 - 80 kHz range. Stimuli were synthesized in Matlab software (Mathworks) and output from a National instruments board (NI-6211, 200 kHz sampling rate) to the ED1 amplifier. Each pure-tone was 500 ms in duration, with 5 ms and 495 ms raised-cosine attack and decay ramps, respectively. The frequency of each pure-tone was randomly selected from 10 equiprobable values (2-45 kHz, 2 tones per octave). Each frequency was repeated 20 times per experiment, with inter-stimulus intervals randomized according at 6, 7, or 8 s. The full stimulus set was presented within approximately 20 minutes.

#### Chronic window implantation

Mice were given an IP injection of dexamethasone (5mg/kg) at least 1 hour prior to surgery to prevent inflammation and edema. Mice were deeply anesthetized using isoflurane (5% induction, 0.5-2% for maintenance) and given a subcutaneous injection of cefazolin (500mg/kg). Internal body temperature was maintained at 37.5 C using a feedback-controlled heating blanket. Scalp fur was trimmed using scissors and any remaining fur was removed using Nair. The scalp was disinfected with alternating swabs of 70% ethanol and betadine. A patch of skin over the temporal bone was removed and the underlying bone cleared of connective tissue using a scalpel. The temporal muscle was detached from the skull, and the skull was cleaned and dried. A thin layer of cyanoacrylate glue (VetBond) was applied to the exposed skull surface and a 3D printed stainless steel head-plate was affixed to the midline of the skull. Dental cement (C&B Metabond) was used to cover the entire head-plate. A circular craniotomy (3 mm diameter) was made over the auditory cortex where the chronic imaging window was implanted. The window was either of a stack of two 3 mm diameter coverslips or a 3.2 mm diameter, 1 mm thick uncoated sapphire window (Edmund Optics), glued with optical adhesive (Norland 61) to a 5 mm diameter coverslip. The space between the glass and the skull was sealed with a silicone elastomer (Kwik-Sil). The edges of the glass and the skull were then sealed with dental cement. Finally, the entire implant except for the imaging window was coated with black dental cement created by mixing methyl methacrylate with iron oxide powder to reduce optical reflections. Meloxicam (0.5mg/kg) was given subcutaneously as a post-operative analgesic.

#### Widefield imaging

To localize primary auditory cortex (A1) via rostro-caudal tonotopic gradients, we performed widefield imaging experiments prior to 2-photon imaging. Awake mice were placed into a 3D-printed plastic tube and head-restraint system. Blue excitation light was shone by an LED (470 nm) through an excitation filter (470 nm) and directed into the cranial window. Emitted fluorescence (F) from neurons in Thy1-GCaMP6s mice was collected through a 4x objective (Thorlabs), passed through a longpass filter (cutoff: 505 nm), followed by a bandpass emission filter (531 nm) attached to a CMOS camera. Images were acquired using ThorCam software (Thorlabs).

After acquiring an image of the cortical surface, the focal plane was advanced to approximately 750 μm below the surface for imaging neural activity. To visualize tonotopy, pure-tones were presented from a free field speaker, as described above. Widefield images were acquired at a 5 Hz rate and 512×512 pixels. Using Matlab software, image sequences for each tone frequency were averaged and background illumination was reduced using a homomorphic contrast filter. For each pixel, ΔF/F traces were calculated by finding the average F taken from the silent baseline period before a pure-tone presentation, subtracting that value from subsequent time-points until 3s after the pure-tone, then dividing all time-points by the baseline F. To visualize auditory responses, we kept traces with ΔF/F within 90% of the maximum response in the pixel-wise grand-average of ΔF/F (i.e., ΔF/F_90_). Pixel-wise tonotopic frequencies were taken as the median frequency of the set of tones corresponding to the ΔF/F_90_ traces.

#### 2-photon imaging

After localizing A1 using widefield imaging, recording sites were selected for 2-photon (2P) imaging in A1 layer 2/3 (L2/3) for each mouse. Our 2P recording sites were chosen at a different region in A1 for each 2P imaging session. Figure 1a illustrates the timeline of our imaging experiments. For each mouse, we began with a pre-injection imaging session (Pre). We then waited an hour before injecting either 2 mg/kg psilocybin (N=6 mice, N=8 experiments) or 0.1 ml saline (N=3 mice, N=6 experiments). The injected mouse was then immediately placed back under the microscope for post-injection imaging (Phase 1), which began within 5 minutes after the injection. With the injected mouse still under the microscope, we then performed a final imaging session (Phase 2).

Our experiments used a scanning microscope (Bergamo II series, Thorlabs) coupled to a pulsed femtosecond 2-photon laser with dispersion precompensation (Coherent Chameleon Discovery NX TPC). The microscope was controlled by ThorImage software. The laser was tuned to λ = 940 nm to excite GCaMP6s. Fluorescence signals were collected through a 16x 0.8 NA microscope objective (Nikon). Emitted photons were directed through a 525 nm (green) bandpass filter onto a GaAsP photomultiplier tube. The field of view was 411 x 411 µm. Imaging frames of 512×512 pixels (0.8 µm per pixel) were acquired at 30 Hz by bidirectional scanning of an 8 kHz resonant scanner. Laser power was set to approximately 70 mW, measured at the objective. During experiments, the objective’s focal plane was lowered into L2/3 (∼100 µm below the surface) before imaging neuronal responses to pure-tones.

After 2P experiments, all images were processed using Matlab. Image motion was corrected using the TurboReg plug-in for MIJI (i.e., FIJI for Matlab). The example 2P fields of view in figure 2a show the averages of registered images for GCaMP6s images across a session. To extract neuronal fluorescence traces, the centers of cell bodies were manually selected, a then a ring-like region of interest (ROI) was cropped around the cell center. Overlapping ROI pixels (due to neighboring neurons) were excluded from analysis. For each labeled neuron, a raw fluorescence signal over time was extracted from somatic ROIs. Pixels within the ROI were averaged to create individual neuron fluorescence traces, _C_(t), for each trial of the experiment. Neuropil fluorescence was estimated for each cellular ROI using an additional ring-shaped ROI, which began 3 pixels from the somatic ROI. Pixels from the new ROI were averaged to obtain neuropil fluorescence traces, _N_(t), for the same time-period as the individual neuron fluorescence traces. Pixels from regions with overlapping neuropil and cellular ROIs were removed from neuropil ROIs. Neuropil-corrected cellular fluorescence was calculated as _C_(t) = _C_(t) – 0.7 _N_(t). Only cells with positive values obtained from averaging _C_(t) across time were kept for analysis, since negative values may indicate neuropil contamination. ΔF/F was calculated from _C_(t), for each neuron, by finding the average F taken from the silent baseline period before a pure-tone presentation, subtracting that value from subsequent time-points until 3s after the pure-tone, then dividing all time-points by the baseline F. ΔF/F was normalized by the standard deviation of response amplitude (σ) for each neuron, yielding the quantity, ΔF/F (σ). Noise correlations were calculated as the correlation coefficient between mean-subtracted neural responses to tones, for each pair of neurons in an experiment.

#### Quantification and statistical analysis

All statistical comparisons were performed using a non-parametric bootstrap t-test with 10000 iterations. All mean values are reported with standard errors of the mean (SEMs).

## SUPPLEMENTAL MATERIALS

**Supplementary Video S1.**
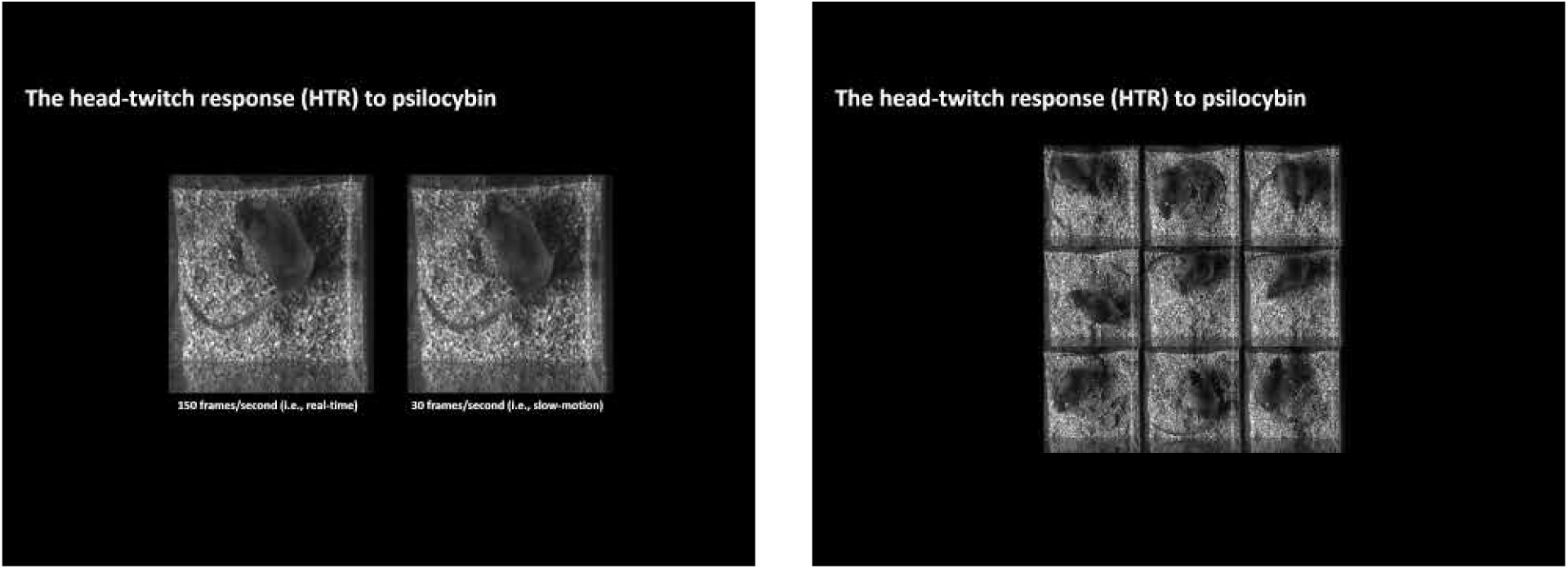
(click the image to access video). Real-time and slow-motion video of head-twitch responses (HTRs). Mice were videoed from above in a clear acrylic box with bedding. Videos were made at 150 frames per second. To better visualize fast mouse movement, videos were analyzed at 30 frames per second to facilitate manual labeling of HTRs. HTRs were reliably observed and labeled after psilocybin injection.

**Supplementary Video S2.**
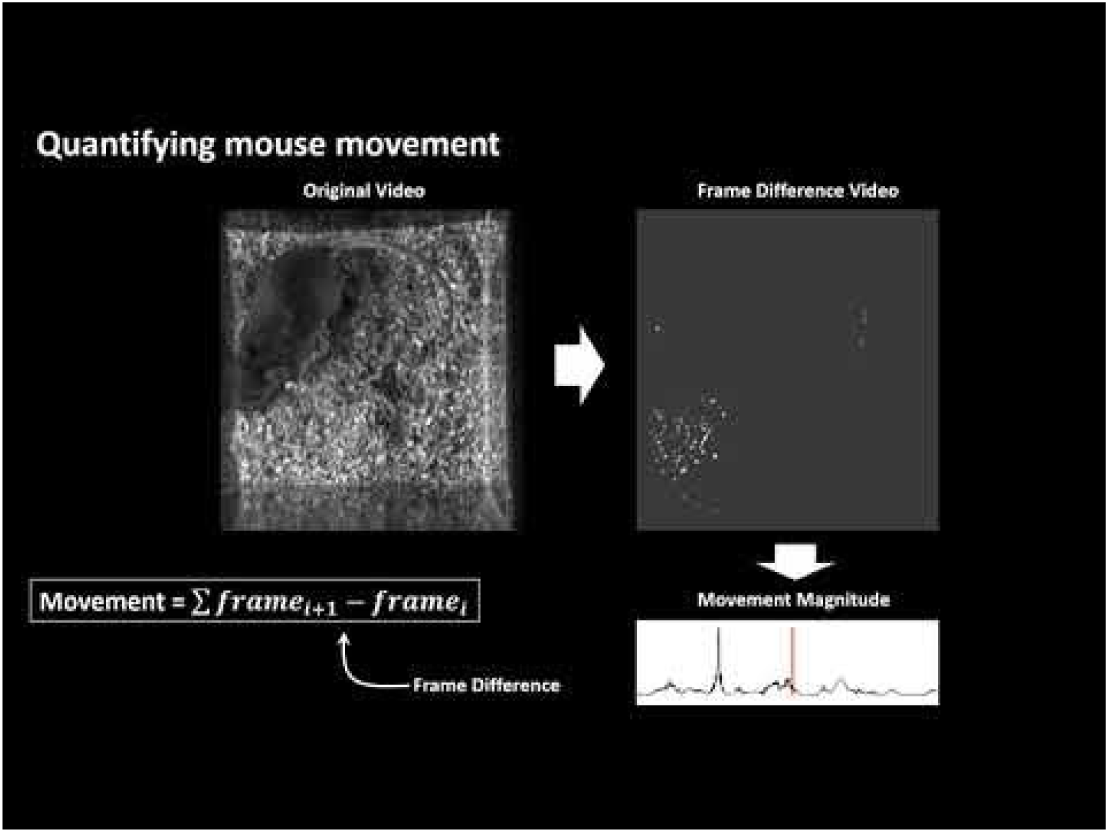
(click the image to access video). Computation of behavioral movement. A frame differencing algorithm was used to quantify mouse movement around the cage (*see* STAR Methods). The video shows how pixel brightness in the frame difference video indicates mouse movement.

## Notes

### Competing Interest Statement

The authors have declared no competing interest.

